# A rapid method to determine the genetic lineage of *Escherichia coli* using open reading frame composition in the shallow sequencing

**DOI:** 10.1101/2024.07.11.601999

**Authors:** Nobuyoshi Yagi, Ayumi Uechi, Itaru Hirai

## Abstract

Determining the genetic background of bacterial isolates and evaluating the genetic relatedness among these isolates in a short time period are important to identify the spreading route(s) in cases of healthcare-associated infections and outbreaks caused by antimicrobial-resistant bacteria. Previously, we proposed a shallow sequencing (Shall-seq) procedure to determine the genetic backgrounds of clinical isolates using the minimum amount of sequence data. However, it took a longer time, such as longer than 10 h, to determine the genetic background of one clinical isolate. In this study, we developed a search procedure using open reading frame (ORF) composition to select the reference genome sequence with the highest matching ratio (>90%), the indicator genome sequence (IGS), for the examined bacterial isolate. Consequently, IGSs were selected for 28 (96.6%) of the examined 29 isolates and selection was performed within 30 min for each bacterial isolate. More importantly, the comparison of IGSs indicated that the IGSs, determined by ORF composition, of the examined bacterial isolates were closely related to the genome sequences determined using the Shall-seq procedure. Taken together, these results suggest that our newly developed search procedure can quickly determine the genetic background of bacterial isolates.

## Introduction

Healthcare-associated infections caused by antimicrobial-resistant bacteria and outbreaks of pathogenic and/or antimicrobial-resistant bacteria occasionally occur. In these cases, determining the genetic background of the causative bacterial isolates and evaluating their genetic relatedness are essential to trace the spreading route of these causative bacteria and to take effective countermeasures.

In our previous study, we established a Shall-seq procedure consisting of nanopore sequencing and data analysis with a minimum amount of sequence data of bacterial isolates[1]. The Shall-seq procedure could be used to select a bacterial genome sequence that was genetically homologous to the sequenced isolate (indicator genome sequence, IGS) from the nucleotide database and detect antimicrobial-resistance genes matched with the isolate’s antimicrobial resistance phenotype. Although the Shall-seq procedure would provide information related to bacterial genetic backgrounds, it took a longer time, such as longer than over 10 h, to select IGS from the nucleotide database. This could be a disadvantage for applying the Shall-seq procedure in clinical practice.

Molecular epidemiology classifying bacterial isolates by patterning the presence or absence of ORFs in their genomes has been proposed [2]. This implies that a homology search using ORF composition could shorten the time required to select IGS for the examined bacterial isolates. In this study, we established a data analysis procedure to select IGS using the minimum sequence data of bacterial isolates obtained from the shall-seq procedure in a shorter time.

## Materials and Methods

### Sequence data

Nanopore sequence data from 29 *Escherichia coli* clinical isolates were obtained from a previous study. The characteristics of the 29 *E. coli* clinical isolates were previously described[1].

### *E. coli* reference genome sequences

A total of 3,186 *E. coli* genome sequences were used, as previously described[1].

### Constructing an *E. coli* ORF database

We annotated 3,186 reference sequences by Prokka [3] using the commands listed in **Table S1**. Consequently, a total of 145,742 sequences annotated as “gene” were collected from the 3,186 reference genome sequences by PanTA [4]. The collected “gene” sequences with sequence identity of 90% or more were sorted into the same ORF using the VSEARCH [5]. Finally, 33,606 “gene” sequences were identified and a database containing these “gene” sequences was designated as an “*E. coli* ORF database”.

### Constructing the ORF datasets of 3,186 reference genome sequences

The ORFs of the 3,186 reference genome sequences were detected using the BLASTN and ORF databases under conditions of over 60% query coverage and over 90% identity. A dataset containing the ORF composition of the reference genome sequences was created and designated as the ORF dataset of the reference genome sequences.

### Selecting indicator genome sequences using the *E. coli* ORF database

The ORFs in the sequence reads of the clinical isolates were detected under conditions where query coverage and percent identity were greater than 90% using KMA [6] and the ORF database. We calculated the logical conjunction between the ORF composition of the clinical isolate and each of the 3,186 reference genome sequences. The matching ratio was then calculated:

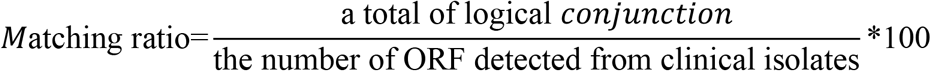

Among the 3,186 reference genome sequences, the reference genome sequence(s) with the highest matching ratio (>90%) was designated as the IGS for clinical isolates.

### Comparison between IGS_SS_ and IGS_ORF_ by constructing phylogenetic tree and drawing a dot plot

To evaluate the difference between the IGS_ORF_ and IGS_SS_, we compared representative 281 *E. coli* genome sequences, containing 23 IGSs determined by ORF composition and Shall-seq procedure. We phylogenetically analyzed these 281 genomes based on the core-genome using Roary[7]. then, phylogenetic tree was constructed by IQTREE2[8] and drawn by R and RStudio. Moreover, we compared IGS_SS_ and IGS_ORF_ and drew dot plots using D-GENIES [9] for clinical isolates wherein the IGS_ORF_ was different from the IGS_SS_.

## Result and Discussion

The ORF composition of the examined *E. coli* clinical isolates was used to search for the genome sequence(s) with the highest matching rate, exceeding 90%, from the ORF dataset of reference genome sequences. Suitable genome sequences were obtained for 28 of the 29 clinical isolates examined. The obtained sequences were designated as indicator genome sequences and IGS_ORF_ of the examined clinical isolates (**Table 1**).

**Table 1.**
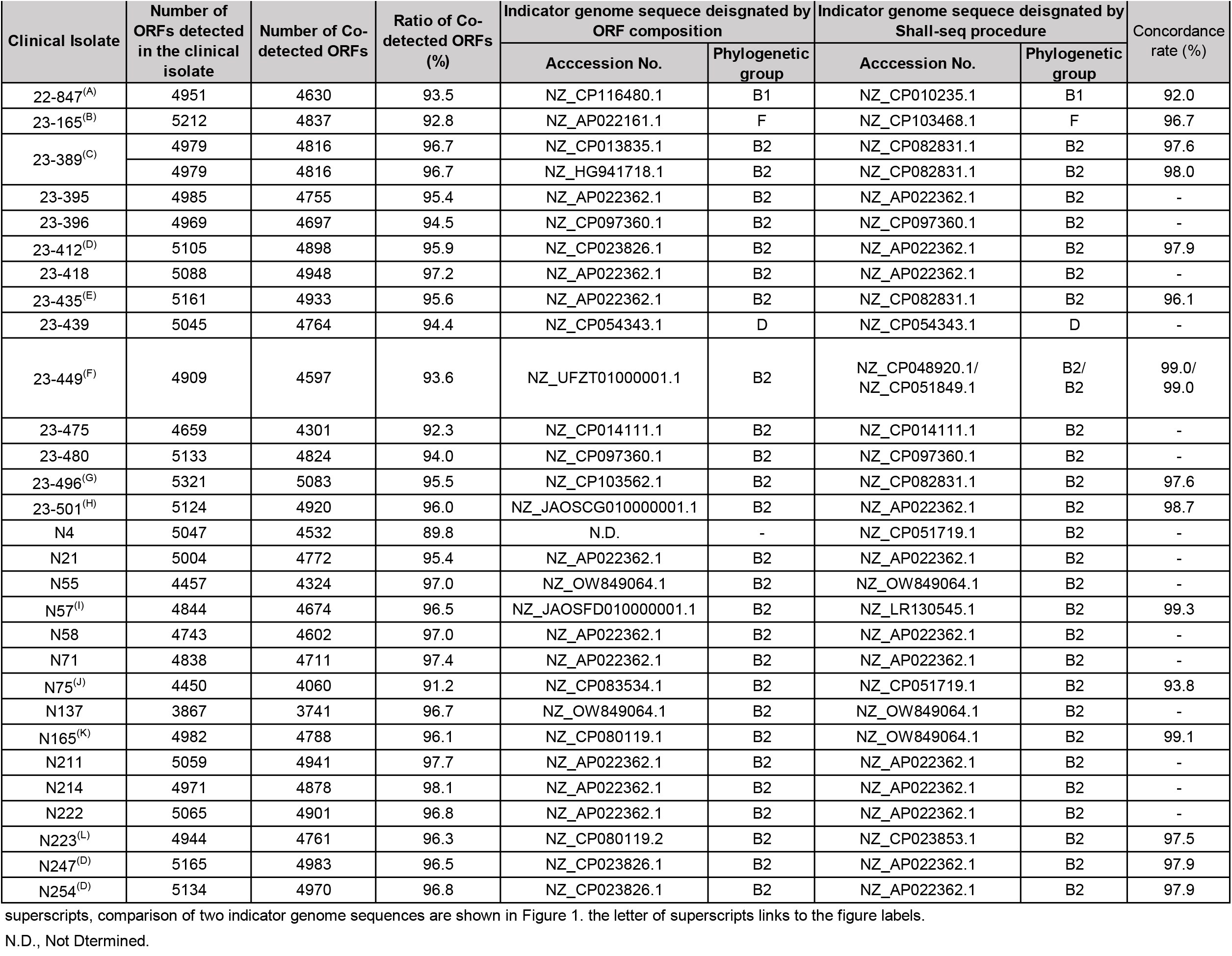
Indicator genome sequences of the clinical isolates.

The IGS_ORF_s differed in 15 (53.6%) of the 28 clinical isolates from the IGS designated by the shall-seq procedure (IGS_SS_), even though there was no discrepancy between them in the phylogenetic groups (**Table 1**). Therefore, we evaluated the difference between IGS_ORF_ and IGS_SS_ by phylogenetic tree (**Fig. 1**). The IGSs determined by ORF composition was closely related to the genome sequences determined using the Shall-seq procedure (**Table 1, Fig. 1**). Moreover, we compared the IGS_ORF_ and IGS_SS_ by using D-genies (**Fig. 2**). Even if there were inversions in certain region of these genomes, the concordance rates between IGS_ORF_ and IGS_SS_ was 92.0% or higher (**Table 1**). Taken together, these results indicated that there was only small difference between IGS_ORF_ and IGS_SS_.

**Fig. 1.**
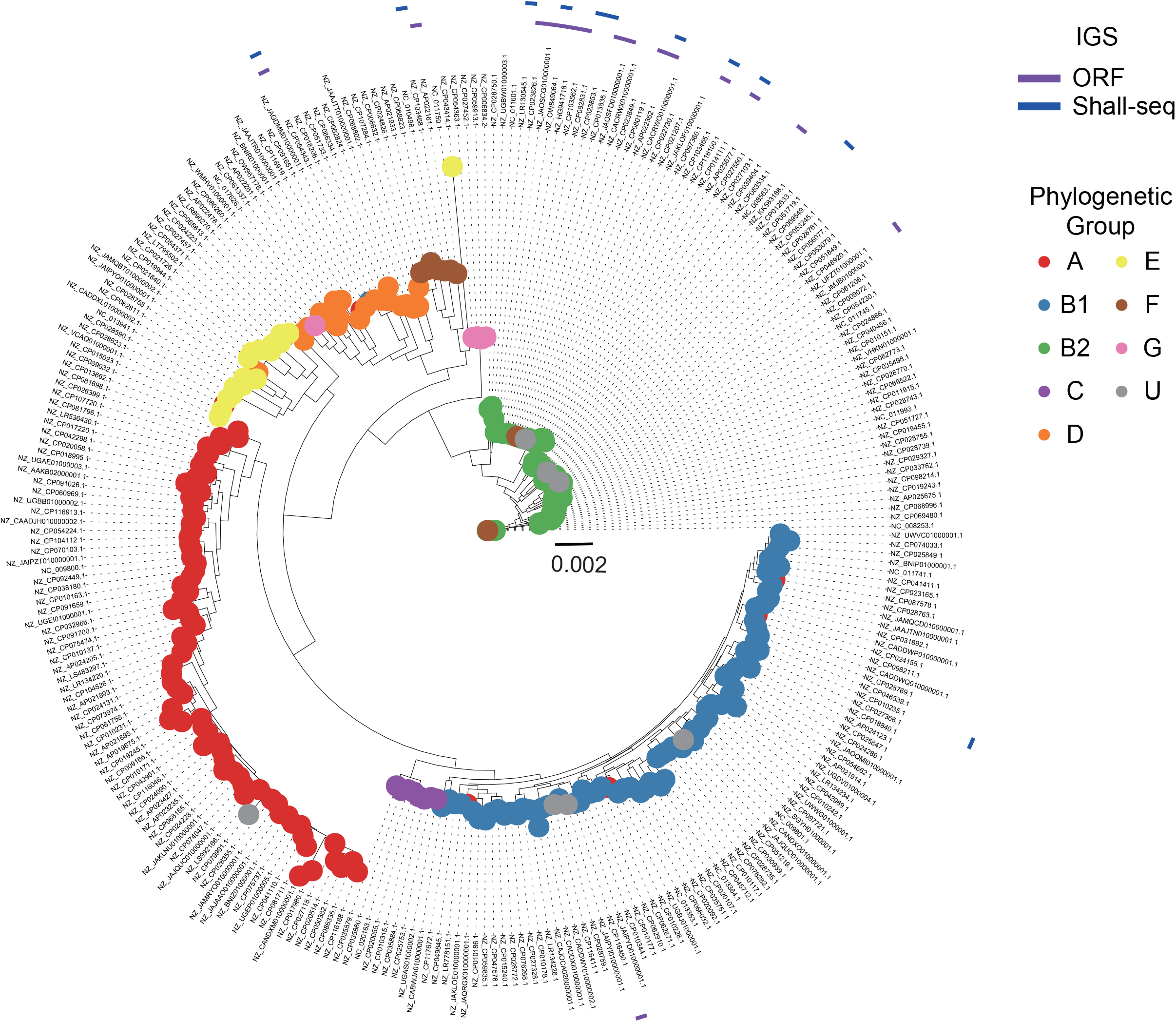
The genetic relationships between IGS_SS_ and IGS_ORF_ by core-genome analysis. The phylogenetic tree for 281 *E. coli* genome sequencies which containing a total of 23 *E. coli* genome sequences determined as IGS_SS_ and IGS_ORF_ was constructed by the Roary. The phylogenetic tree was constructed by 1474 genes detected as core gene with the Roary and IQtree2. The circle plot at the end of the branch means the phylogenetic group of each *E. coli* reference genomes. The heat map at the outside of phylogenetic tree means the *E. coli* genomes detected as IGS by ORF composition (purple) and/or Shall-seq procedure (blue).

**Fig. 2.**
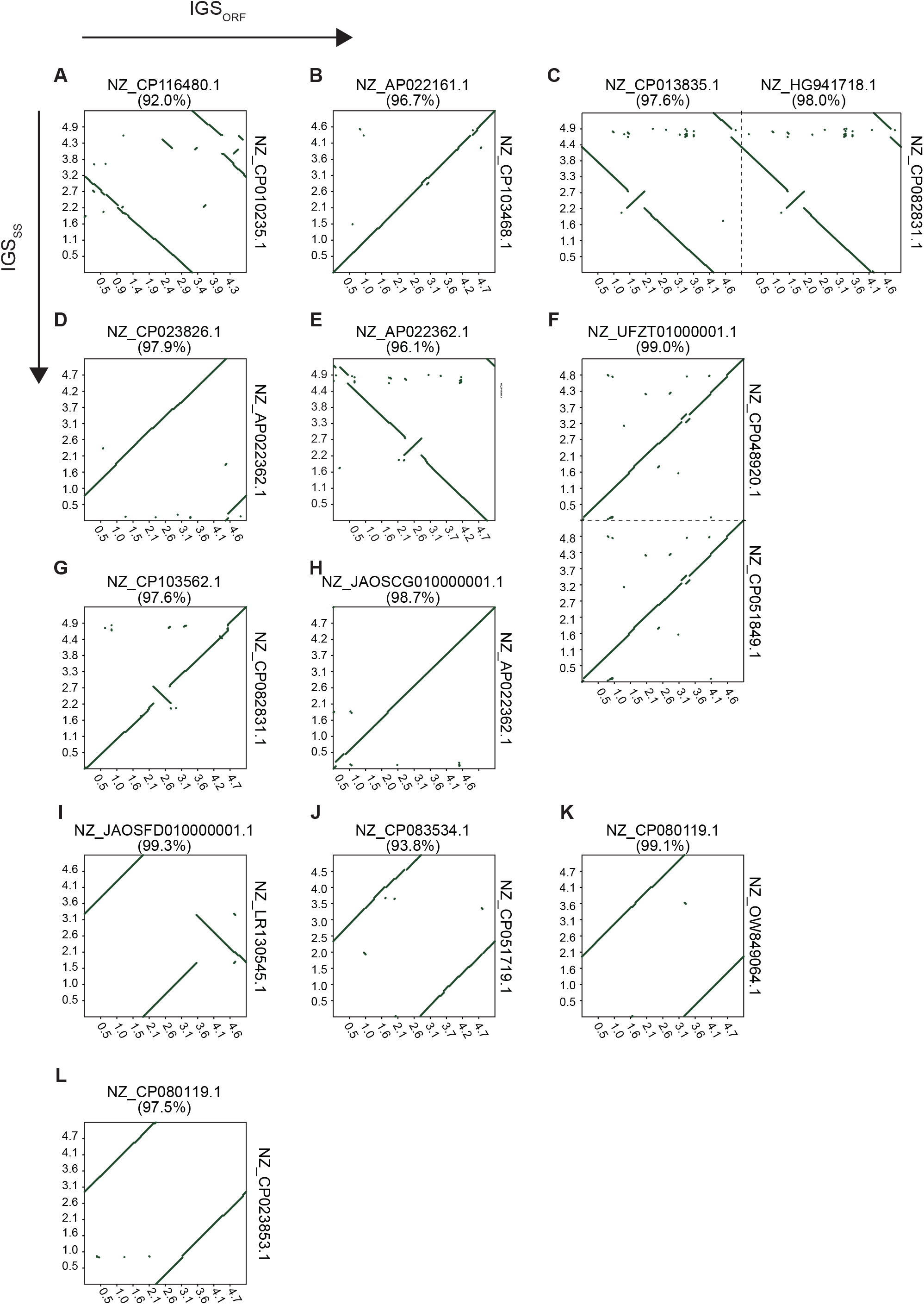
Comparison between IGS_ORF_ and IGS_SS_. In 15 clinical isolates, different reference genome sequences were selected between IGS_ORF_ and IGS_SS_, and IGS_ORF_s (horizontal axis) and compared with IGS_SS_s (vertical axis) by drawing dot plots using D-Genies. Each dot represents > 75% nucleotide identity. Within the plot, diagonal lines indicate homologous regions between the two indicator genomes; regions lacking such diagonals comprise each indicator’s genome-specific regions. Figure labels are linked to the superscripts in **Table 1**.

It took 10 h per isolate to determine IGS_SS_ when the shall-seq procedure was performed. The application of the ORF composition reduced the time from 10 h to 30 min per isolate. It was suggested that IGS_ORF_ would be obtained within one hour after obtaining sequence reads of the Nanopore sequencing even considering the time required for quality check of output sequence reads (approximately 15 min).

As mentioned above, our data indicated that the IGS_ORF_ was obtained by a combination of the shall-seq procedure and a search using the ORF composition. Many bacterial isolates should be analyzed and genetic relatedness among them should be evaluated as soon as possible when healthcare-associated infections or outbreaks of pathogenic bacteria occur. In this regard, the genetic relatedness among bacterial isolates can be evaluated and visualized by drawing a phylogenetic tree using IGS_ORF_. Until recently, higher error rates were reported as a disadvantage of nanopore sequencing[10]. However, the accuracy of nanopore sequencing has recently improved [11]. Therefore, the improved accuracy of nanopore sequencing might encourage the application of the Shall-Seq procedure and searches using ORF composition.

## Data availability

The sequence reads used in this study are available under BioProject accession number PRJDB17430.

